# Perception of near-threshold visual stimuli is influenced by pre-stimulus alpha-band amplitude but not by alpha phase

**DOI:** 10.1101/2023.03.14.532551

**Authors:** María Melcón, Enrique Stern, Dominique Kessel, Lydia Arana, Claudia Poch, Pablo Campo, Almudena Capilla

## Abstract

Ongoing brain activity preceding visual stimulation has been suggested to shape conscious perception. The underlying mechanisms are still under debate, although alpha oscillations have been pointed out as the main explanatory candidate. According to the pulsed-inhibition framework, bouts of functional inhibition arise in each alpha cycle, allowing information to be processed in a pulsatile manner. Consequently, it has been hypothesized that perceptual outcome can be influenced by the specific phase of alpha oscillations prior to the stimulus onset, although empirical findings are controversial. In this study, we aimed to shed light on the role of pre-stimulus alpha oscillations in visual perception. To this end, we recorded electroencephalographic (EEG) activity while participants performed three near-threshold visual detection tasks with different attentional involvement: a no-cue task, a non-informative cue task (50% cue validity), and an informative cue task (100% cue validity). Cluster-based permutation statistics were complemented with Bayesian analyses to test the effect of pre-stimulus oscillatory amplitude and phase on visual awareness. We additionally examined whether these effects differed on trials with low and high oscillatory amplitude, as expected from the pulsed-inhibition theory. Our results show a clear effect of pre-stimulus alpha amplitude on conscious perception, but only when alpha fluctuated spontaneously and was not modulated by attention, supporting the notion that alpha-band power indexes neural excitability. In contrast, we did not find any evidence that pre-stimulus alpha phase influences the perceptual outcome, not even when differentiating between low and high amplitude trials. Furthermore, Bayesian analysis provided moderate evidence in favor of the absence of phase effects. Taken together, our results challenge the central theoretical predictions of the pulsed-inhibition framework, at least for the particular experimental conditions used here.

## Introduction

Perception is not simply the passive reception of stimuli from the physical environment. Instead, it is the outcome of the interaction between incoming sensory stimulation and the current state of the brain (Arieli et al., 1996). Indeed, the influence of ongoing brain activity is particularly evident when identical stimulation yields different perceptual outcomes, as shown by a growing number of studies using masked stimuli (Benwell et al., 2022; Hanslmayr et al., 2007; Limbach & Corballis, 2016; Mathewson et al., 2009; Ruzzoli et al., 2019) as well as near-threshold stimuli detection tasks (Benwell et al., 2017; Busch et al., 2009; Busch & VanRullen, 2010; Capilla et al., 2014; Chaumon & Busch, 2014; Harris et al., 2018; Iemi et al., 2017; Wyart & Tallon-Baudry, 2008; Zazio et al., 2021). Similarly, the perception of phosphenes induced by transcranial magnetic stimulation (TMS) has also been demonstrated to depend on the state of neural excitability (Dugue et al., 2011; Fakche et al., 2022; Romei et al., 2008; Samaha et al., 2017).

Ongoing brain oscillations reflect rhythmic fluctuations between high and low excitability states (Bishop, 1932; Buzsáki et al., 2012). Critically, neural oscillations in the alpha frequency range (8-12 Hz) have been proposed to modulate the excitation/inhibition balance of cortical neural assemblies in a pulsatile manner (Klimesch et al., 2007; Mathewson et al., 2011; Mazaheri & Jensen, 2010). According to this framework, bouts of functional inhibition arise every 100 ms, rhythmically blocking and opening windows of opportunity for information processing in each alpha cycle. Interestingly, it has been argued that alpha-band amplitude controls the length of the time window of processing (i.e., the duty-cycle). Thus, a low alpha power, reflecting a state of high excitability, would provide long time windows of processing and, consequently, improved perceptual performance. In contrast, an increase in alpha amplitude would lead to short duty-cycles concentrated at the optimal alpha phase for visual processing; while neural processing would be inhibited at the opposite phase (Jensen et al., 2012; Mathewson et al., 2011; Mazaheri & Jensen, 2010).

The pulsed-inhibition view of alpha oscillations has received partial support from experimental work. Whereas most studies agree on the negative relationship between alpha-band amplitude during the pre-stimulus time interval and visual awareness (Benwell et al., 2017, 2022; Busch et al., 2009; Busch & VanRullen, 2010; Capilla et al., 2014; Chaumon & Busch, 2014; Fakche et al., 2022; Iemi et al., 2017; Limbach & Corballis, 2016; Mathewson et al., 2009; Romei et al., 2008; Ruzzoli et al., 2019; Samaha et al., 2017; Zazio et al., 2021), the role of pre-stimulus alpha phase on perception remains highly controversial. The seminal studies of Busch et al. (2009) and Mathewson et al. (2009) found that the detection of near-threshold or masked stimuli was facilitated if alpha-band oscillations were at a specific phase prior to the stimulus onset. Since then, the effect of alpha phase on perception has been successfully replicated in some studies (Busch & VanRullen, 2010; Dugue et al., 2011; Fakche et al., 2022; Harris et al., 2018; Samaha et al., 2017; Zazio et al., 2021), although others have failed to find any phase effect (Benwell et al., 2017, 2022; Chaumon & Busch, 2014; Ruzzoli et al., 2019), including two pre-registered reports (Ruzzoli et al., 2019; Vigué-Guix et al., 2022).

Unravelling this discrepancy is crucial, as alpha phase lies at the heart of the theoretical accounts on the functional role of the alpha rhythm in perception. Inconsistencies in the literature might be due to the use of different experimental designs and analysis parameters, as well as to a potential interaction between alpha amplitude and phase. First, different studies have made use of diverse experimental paradigms, in particular, regarding the involvement of anticipatory attention. Some studies have investigated the effect of the pre-stimulus phase of the spontaneously fluctuating alpha rhythm (e.g., Busch et al., 2009; Hanslmayr et al., 2005), while others have modulated stimulus expectation and pre-stimulus alpha activity by anticipatory cues with different validity percentages (Harris et al., 2018; Wyart & Tallon-Baudry, 2008). Second, different analysis choices (i.e., electrodes and frequency ranges) might also explain the discrepancies found between studies. Whereas amplitude effects are relatively robust and often restricted to parieto-occipital electrodes in the alpha band (Chaumon & Busch, 2014; Hanslmayr et al., 2007; Limbach & Corballis, 2016; Ruzzoli et al., 2019; Samaha et al., 2017; van Dijk et al., 2008; Zazio et al., 2021), phase effects are rather heterogeneous. They have been found at fronto-central (Han & VanRullen, 2017; Samaha et al., 2017; Zazio et al., 2021) as well as parieto-occipital electrodes (Han & VanRullen, 2017; Harris et al., 2018; Samaha et al., 2015, 2017; Sherman et al., 2016) over a wide range of oscillatory frequencies, from theta (Han & VanRullen, 2017; Harris et al., 2018; Wutz et al., 2016) to alpha (Harris et al., 2018; Samaha et al., 2015, 2017; Sherman et al., 2016) and beta (Han & VanRullen, 2017). Finally, as predicted by the pulsed-inhibition framework, alpha phase would only be expected to influence subsequent perception when alpha-band power is high (Mathewson et al., 2011; Mazaheri & Jensen, 2010). However, most previous work has analyzed all high and low alpha amplitude trials together, which may partially explain why alpha phase results are not consistent.

In this study, we aimed to contribute to the current controversy on the role of pre-stimulus alpha-band oscillations in visual perception. To this end, we designed a near-threshold detection task, in which visual stimuli appeared to the left or to the right of a fixation point. We investigated the influence of attention on subsequent perception manipulating attentional engagement by means of an anticipatory cue. Thus, all participants performed three tasks: a no-cue task, an informative cue task (spatial cue 100% valid) and a non-informative cue task as a control (spatial cue 50% valid). For all three tasks, we followed a data-driven approach and avoided any a priori selection of analysis parameters to examine the effect of pre-stimulus oscillatory amplitude and phase, as well as the differential effect of phase in trials with high and low amplitude. Finally, in addition to traditional frequentist statistics, we applied Bayesian analysis to determine which of the two scenarios gathered more evidence: either the existence or the absence of alpha amplitude and phase effects.

## Materials and methods

### Participants

Sample size was estimated using G*power version 3.1.9.7 (Faul et al., 2007) for a medium effect size of 0.5, an α-value of 0.05, and an estimated power of 0.80. We obtained a minimum sample size of 34 participants. However, in anticipation that some of them could be excluded from the analysis due to artifacts in the EEG signal, we decided to recruit 2 additional participants. Thus, thirty-six healthy graduate students (21.0 ± 6.2 years old, mean ± SD, 27 females, 30 right-handed) from the Universidad Autónoma de Madrid with normal or corrected-to-normal vision took part in this experiment. The participation was voluntary and they all provided prior informed consent. The experimental protocol complied with the declaration of Helsinki and was approved by the Ethics committee of the Universidad Autónoma de Madrid.

### Stimuli

Stimulation consisted of vertically and horizontally oriented Gabor patches. They were generated online using MATLAB (R2021b) and presented at two locations, 5° below and 5° to the right or the left from the fixation cross. Gratings were presented near the threshold of conscious perception. To mitigate the effect of factors such as fatigue or habituation over the course of the experiment and achieve the best adjustment, the threshold was estimated using an online adaptive procedure (Dixon & Mood, 1948). Thus, the contrast of the Gabor was dynamically calculated, with each trial being determined by the responses to the previous trials based on an adaptation of the transformed up-down staircase (Levitt, 1971), as explained in the following section.

### Experimental procedure

Psychtoolbox was used to control stimulus presentation and behavioral data collection (v3.0; Brainard, 1997). Stimulation was presented on a 16-inch monitor, while participants were comfortably seated 65 cm away from the screen in a dimly lit, sound-attenuated, and electromagnetically shielded room. To improve timing control, stimulus onset was monitored by using a photodiode. Participants completed three near-threshold visual stimulus detection tasks with different degree of attentional involvement: no-cue, non-informative cue (50% validity), and informative cue (100% validity) (see Figure 1).

**Figure 1.**
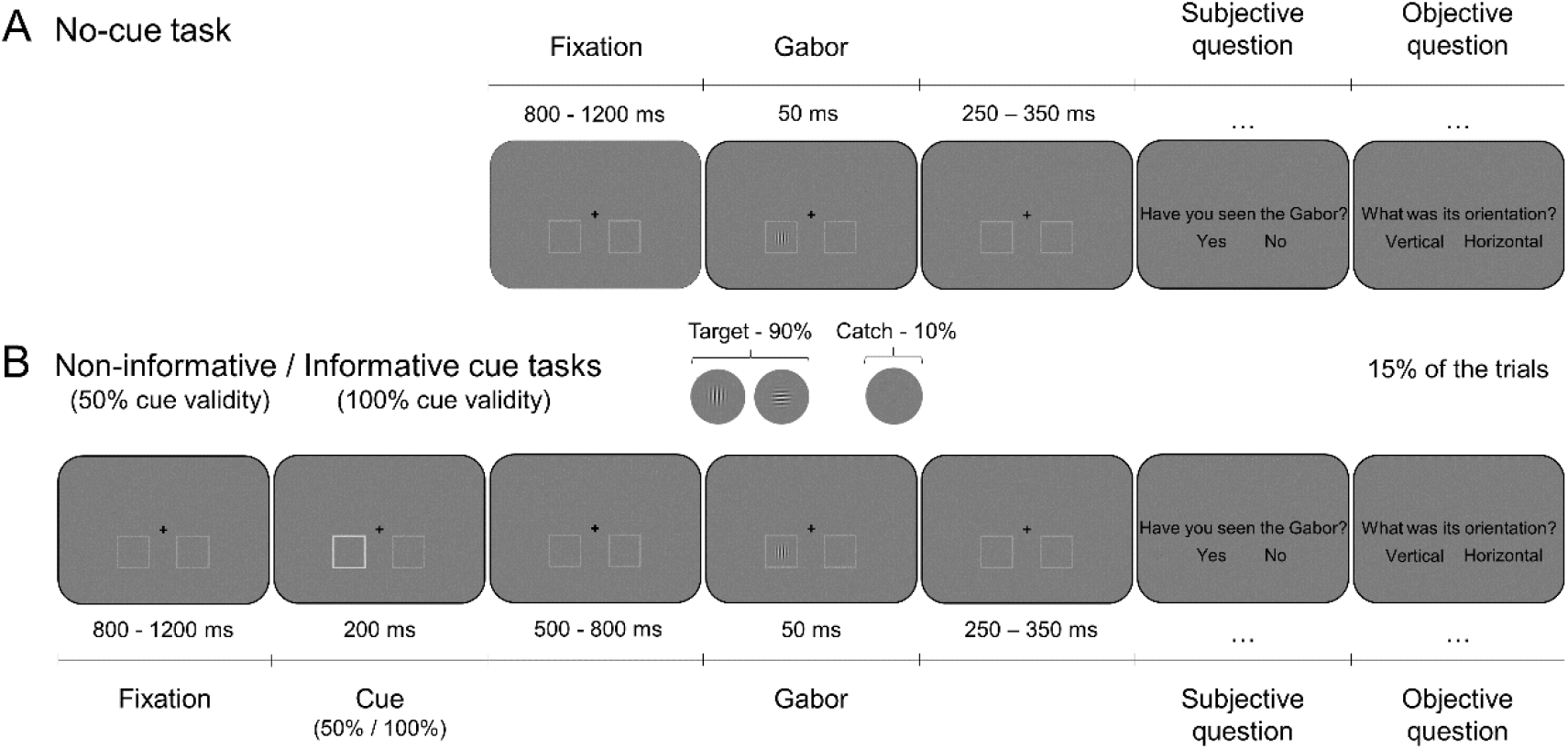
Experimental task. Participants conducted three near-threshold visual detection tasks with different attentional involvement. Stimulus contrast was calibrated online on a trial-by-trial basis. Participants were asked to report the appearance of the Gabor stimulus (subjective question), and also to indicate the Gabor orientation in a small percentage of trials (objective question). **A.** No-cue task. No cue preceded the Gabor stimulus. **B.** Non-informative and informative cue tasks. A cue was presented before the Gabor onset with 50% and 100% validity for each task, respectively.

In the no-cue task, each trial started with a fixation cross on a dark grey background and two boxes outlined in light grey, placed at the lower left and right side of the fixation cross (Figure 1A). After a random delay of 800-1200 ms, a grating was presented at peri-threshold contrast during 50 ms. Gabor stimuli were equally likely to appear at either the left or the right location and to be either vertically or horizontally oriented. As a control, no stimulus was presented on 10% of trials (catch trials). After a variable period of 250-350 ms, behavioral responses were collected. First, participants had to answer a subjective question, reporting whether they had seen or not any Gabor stimulus (i.e., response was yes/no). Then, in 15% of randomly selected trials, they were additionally asked to indicate whether the orientation of the Gabor was horizontal or vertical (objective question). Both response displays consisted of a question and the two response alternatives (yes/no, horizontal/vertical) located on the right and the left. Participants provided their responses by pressing a key on a numerical keyboard with the corresponding right or left thumb. The position of the response alternatives varied randomly in each trial to avoid motor preparation. Participants had unlimited time to provide their responses and were suggested to blink during this period. This task comprised a total number of 400 trials (180 Gabor stimuli in each visual hemifield and 40 catch trials), with self-paced breaks every 100 trials to avoid fatigue.

Both the informative and the non-informative cue tasks had the same timing and events as the task described above, except for a cue and an orienting period after the fixation (Figure 1B). The cue consisted of one of the two boxes turning white for 200 ms (equal probability for left and right boxes). After a variable interval of 500-800 ms, the Gabor was presented in either the left or the right box. In the non-informative cue task, the cue indicated with a validity of 50% the location of the upcoming Gabor, whereas in the informative cue task, the cue was 100% valid. The total number of trials in each task was also 400, including 40 catch trials and self-paced breaks.

To increase the variability in stimulus contrast, this varied from trial to trial by adding a random quantity to the threshold contrast. The threshold was dynamically adjusted for each participant, task, and visual field in two steps: a pre-task calibration and an in-task calibration. Both steps were based on the transformed up-down staircase algorithm (Levitt, 1971).

The goal of the pre-task calibration was to find an initial perceptual threshold. Here, the Gabor was presented with 10 pre-set contrast values for each hemifield. The pre-task calibration finished when the percentage of seen responses was in the range between 30% and 70%. If performance was out of that range, participants were required to conduct a new calibration series where the contrasts were adjusted according to their responses (decreasing the values for <30% unseen responses and increasing the values for >70% unseen responses). This calibration not only established the initial threshold to start the task, but also served as a practice session.

The in-task calibration procedure was similar to the previous one. However, to avoid the calibration being stuck at an erroneous threshold, we calculated the stimulus contrast on a trial-by-trial basis rather than for the whole series. Stimulus contrast was initially set to a value around the previously obtained threshold. This value changed dynamically during the task based on the responses to a sequence of immediately preceding trials. The length of this sequence was randomly determined on each trial, from 1 to 4. If the sequence comprised only seen responses, the contrast of the upcoming stimulus decreased, while a succession of unseen responses was required for the stimulus contrast to increase.

Each task started with the presentation of the instructions to the participants. They were informed about the brief appearance of the Gabor, the variability in contrast levels, the presence or absence of a cue, and whether the cue provided information about the upcoming stimulus. Participants were asked to maintain their gaze on the fixation cross throughout the experiment and were instructed to respond to both the subjective and the objective tasks as quickly and accurately as possible. The three tasks were presented in random order. Participants were encouraged to remain still and relaxed during the whole experiment, which lasted approximately 60 minutes.

### Behavioral analysis

Behavioral analyses were carried out to test (1) the Gabor contrast, since it was expected to differ according to both attention (no-cue, non-informative cue, and informative cue) and awareness (seen vs. unseen), and (2) that participants were engaged in the task, meaning that their performance in the objective task was above chance level. Statistical analyses were conducted using the SPSS 16.0 software package.

For the first analysis, we conducted a 3×2×2 repeated measures analysis of variance (ANOVA) for Gabor contrast. Specifically, we tested the effect of Attention (no-cue, non-informative cue, informative cue), Awareness (seen, unseen), and Hemifield (left, right). We considered Greenhouse-Geisser correction in case of non-sphericity. To detect specific differences between conditions, post-hoc pairwise t-tests were conducted using Bonferroni correction for multiple comparisons. Effect sizes were estimated using the partial eta-squared (η_p_^2^) method. Only data from twenty-five participants were included in this ANOVA, those whose EEG data were analyzed in the three tasks.

Secondly, we calculated the percentage of correct responses in the objective forced-choice question, when participants reported the Gabor orientation. We then performed t-tests vs. 0.5 (i.e., chance level) independently for each task (i.e., no-cue, non-informative cue, and informative cue). Since statistical analyses were conducted separately for the three tasks, we included the same participants as in the EEG analysis for each of them.

### EEG recording

Data were acquired using a BioSemi ActiveTwo system with 128 EEG channels, four external electrodes (placed below and above one eye, and lateral to both eyes), and another one as a potential reference on the nose-tip. Offsets of the active electrodes were kept below 25-30 mV. Data were digitized at a 1024 Hz sampling rate and low-pass filtered online at 205 Hz.

### EEG data analysis

EEG data were analyzed using the Fieldtrip toolbox (Oostenveld et al., 2011) and in-house MATLAB code. The aim of these analyses was to detect pre-stimulus oscillatory amplitude and phase differences between seen and unseen trials. The pipeline was applied independently for each task: no-cue, non-informative cue, and informative cue. After data preprocessing, the EEG signal was time-frequency decomposed and amplitude effects were tested by means of a cluster-based permutation statistical analysis. Next, we investigated whether seen and unseen trials exhibited opposite phase clustering by computing the Phase Opposition Sum (POS) (VanRullen, 2016a). Phase effects were also tested by a cluster-based permutation approach, first for all trials together, and then separately for high and low amplitude trials.

#### Preprocessing

We segmented the continuous EEG signal into 3800 ms epochs, starting 2000 ms before the Gabor onset. These long epochs aimed to avoid edge artifacts in subsequent time-frequency analyses. An artifact rejection procedure was then applied in three steps. First, we visually inspected the signal to discard trials contaminated by eye or cable movements, swallowing, or muscular activity. Participants with more than 45% of trials contaminated with ocular artifacts were not included in subsequent analyses (5 participants in the no-cue task, 6 participants in the non-informative cue task, and 5 participants in the informative cue task). Then, we removed any remaining artifacts by means of Independent Component Analysis (ICA, ‘runica’ algorithm). Finally, noisy electrodes were interpolated based on the signal from adjacent electrodes (mean ± SD of interpolated electrodes: 1.9 ± 2.1 for the no-cue task, 2.5 ± 2.9 for the non-informative cue task, and 3.1 ± 2.7 for the informative cue task). Finally, data were downsampled to 512 Hz, linearly detrended, baseline corrected (−500 to 0 ms), and re-referenced to the common average.

Data were then split into four experimental conditions based on Gabor location (left/right) and awareness (seen/unseen). Importantly, since Gabor contrast was dynamically changing throughout the experiment, we additionally controlled that the contrast of the analyzed seen and unseen trials did not differ. This was achieved by iteratively discarding the trials with the highest contrast from the seen condition and the trials with the lowest contrast from the unseen condition, until Gabor contrast did not significantly differ between them (p > .05). Thus, the final number of analyzed trials for the no-cue task was 307 ± 35, 290 ± 38 trials for the non-informative cue task, and 256 ± 47 trials for the informative cue task (mean ± SD).

Finally, one participant in the no-cue task was discarded because contrast differences between seen and unseen trials remained significant after applying the above procedure. In addition, behavioral data from one participant in the no-cue task and two participants in the non-informative cue task could not be saved due to technical reasons, so they were also excluded from subsequent analyses. In sum, the number of participants analyzed was 29 in the no-cue task, 28 in the non-informative cue task and 31 in the informative cue task.

#### Time-frequency analysis

Time-frequency decomposition was calculated using a sliding window Fourier transform with a Hanning taper in steps of 20 ms. This analysis was conducted on trials of 3800 ms duration (starting 2000 ms before the Gabor onset). We analyzed 15 logarithmically spaced frequencies from 2 to 33 Hz. The length of the sliding window was frequency-dependent, with the number of cycles increasing logarithmically from 2 to 7 cycles.

Then, to estimate the oscillatory activity contralateral and ipsilateral to stimulus location, we transformed right-stimulus trials into a mirrored version. Hence, in both conditions (left and right Gabor stimuli), ipsilateral activity is always represented in left channels, whereas contralateral activity is displayed in right channels. Left and right stimulus conditions remained collapsed for all subsequent analyses. Finally, we set out to test whether pre-stimulus oscillatory amplitude differed between seen and unseen trials in each task (no-cue, non-informative cue, and informative cue). Thus, we first computed the within-participant average time-frequency amplitude separately for seen and unseen trials. Then, we applied a dependent-samples cluster-based permutation test on the three-dimensional (channel-frequency-time) data, which accounted for the multiple comparisons problem (Maris & Oostenveld, 2007). In brief, adjacent electrodes, frequency bins, and time points with uncorrected p-values below 0.05 were grouped into positive and negative clusters (two-tailed t-test). The minimum number of neighbor channels required to be included in a cluster was 4, and the weighted cluster mass was selected as the cluster statistic, since this combines cluster size and intensity (Hayasaka & Nichols, 2004). The significance probability of the cluster statistic was evaluated by means of a permutation test. Data from both conditions (seen and unseen trials) were grouped and randomly assigned to two subsets, from which the maximum cluster-level statistic was computed and extracted. For each task and participant, this procedure was repeated 10,000 times in order to build the null distribution approximated by the Monte-Carlo estimate. The p-values were then computed as the proportion of permutations above the observed cluster-level statistic (α critical value = 0.025, two tails).

#### Oscillatory phase analysis

In addition, we aimed to test whether pre-stimulus oscillatory phase was different in subsequently seen and unseen trials. To this end, we computed the Phase Opposition Sum (POS) (VanRullen, 2016a), which is based on the comparison of the inter-trial phase coherence (ITC; i.e., phase consistency) across trials between conditions. More specifically, POS is calculated according to the following formula:

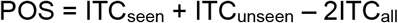

where ITC_seen_ and ITC_unseen_ represent the ITC of each condition and ITC_all_ is the overall ITC of all trials together, which serves as a baseline. This procedure yields values close to 0 when the two conditions do not have a strong ITC and also when their phases are clustered at approximately the same phase angles. On the contrary, a POS value of 1 indicates a complete opposition between the phases of the two conditions. It is important to highlight that POS is a pure phase measure that is not influenced by amplitude differences (Fakche et al., 2022).

To statistically test POS values, we used the same cluster-based permutation approach as for amplitude. However, since POS is obtained from the comparison between the two conditions (seen and unseen trials), we first estimated a null POS per participant. Thus, we could apply the cluster-based permutation analysis to test for differences between empirical (POS_seen-unseen_) and null POS values. The null POS distribution was estimated by randomly drawing seen and unseen trials over 10,000 repetitions and computing ITC and POS in each of them. Eventually, the POS values of this null distribution were averaged resulting in a POS value per participant, electrode, frequency, and time point. We then applied a non-parametric cluster-based permutation analysis to test for differences between empirical and null POS values.

Finally, since the pulsed-inhibition hypothesis (Klimesch et al., 2007; Mathewson et al., 2011; Mazaheri & Jensen, 2010) predicts that phase effects are more prominent when alpha amplitude is high, we additionally repeated the above procedure separately for high and low oscillatory amplitude. To avoid a priori selection of a pre-stimulus time-frequency window to estimate amplitude, we sorted and binned each electrode-frequency-time bin into two groups based on a median-split of the amplitude values.

### Ad-hoc statistical analysis

Given the absence of any significant pre-stimulus phase opposition effect between seen and unseen trials, we decided to conduct additional ad-hoc statistical analyses to test whether any weak effect might emerge under largely liberal statistical assumptions. Thus, we computed the grand-average of the difference between empirical and null POS values, collapsed across electrodes, and selected ad-hoc the time-frequency interval with the highest values. We then performed a paired t-test between empirical and null POS values averaged over the time-frequency window of interest. This analysis was also performed separately for trials with high and low pre-stimulus alpha amplitude. It is important to highlight that the results of this type of analysis are not valid from a statistical point of view, as it is a clear example of “double dipping” (i.e., same data are used for selection and selective analysis; see Kriegeskorte et al., 2009) and multiple comparisons are not adequately controlled for (Maris & Oostenveld, 2007). Nevertheless, we believe that it can provide valuable information by checking for any potential weak phase effects that do not exceed statistical thresholds when appropriate statistical controls are applied.

### Bayesian analysis

Finally, since frequentist statistical tests can only provide evidence against but not in favor of the null hypothesis (H0), we additionally conducted Bayesian analyses (Dienes, 2014). These aimed to test whether there was evidence for the presence/absence of effects in both pre-stimulus oscillatory amplitude and phase between seen and unseen trials. First, clusters obtained in the previous permutation analyses, specifically from the no-cue and the non-informative cue tasks, were now used to extract a mask of common electrodes, frequency bins, and time points to be tested with Bayesian statistics. In addition, since electrodes contributed differently to the cluster, a weight was assigned based on the number of times each electrode was present across time. We thus obtained one amplitude and one POS value per participant and task, resulting from the weighted average across electrodes, frequency bins, and time points within the mask.

Then, we employed JASP 0.17.1 (JASP Team, 2023) to calculate BF_10_, where a value higher than 3 indicates substantial evidence in favor of the alternative (H1) over the null (H0) hypothesis, while values lower than 1/3 supports the H0 against the H1 (Jeffreys, 1998; see also Dienes, 2014). H1 was modelled separately for amplitude and phase using a Cauchy distribution centered on zero and with scale factors estimated from standardized effect sizes (Cohen’s d) reported in previous related studies. Since we expected a similar attentional modulation in the non-informative cue and in the no-cue task, we calculated a common prior for both tasks based on the effect sizes of studies using either a 50% valid cue (Busch & VanRullen, 2010) or no cue (Chaumon & Busch, 2014; Mathewson et al., 2009). The resulting priors were 0.91 for amplitude (Busch & VanRullen, 2010; Chaumon & Busch, 2014) and 0.85 for phase (Mathewson et al., 2009). Regarding the informative cue task, to the best of our knowledge, experiments studying pre-stimulus alpha amplitude have not found any statistical differences between seen and unseen trials, so we decided to employ a prior width of 0.31, corresponding to a small effect size of 0.1 (Harris et al., 2018; Milton & Pleydell-Pearce, 2016; Wyart & Tallon-Baudry, 2008). Finally, we computed the scale factor for the phase analysis based on Milton et al. (2016). We could not find any studies that assessed pre-stimulus alpha phase effects while modulating attention by a 100% valid cue. Hence, we selected the above study for employing the highest cue validity (75%) compared to similar research. The resulting prior width for the phase analysis of the informative cue task was, therefore, 0.75.

## Results

### Gabor contrast was modulated by attention

The first aim of the behavioral analyses was to compare the Gabor contrast between seen and unseen trials. A 3×2×2 ANOVA performed over Gabor contrast with the factors Attention, Awareness, and Hemifield revealed significant main effects of Attention (F_2,48_ = 6.34, p = 0.004, η_p_^2^ = 0.21) and Awareness (F_1,24_ = 332.70, p < 0.001, η_p_^2^ = 0.93), and no interaction effects. Specifically, for Attention, Gabor contrast was higher for both the no-cue and the non-informative cue tasks compared with the informative cue task (Figure 2A). Regarding Awareness, Gabors reported as seen had a higher contrast than those reported as unseen, as expected (Figure 2B). Notice that contrast differences were controlled in further EEG analyses by iteratively discarding the trials with highest/lowest contrast from the seen/unseen condition, respectively, until they did not differ.

**Figure 2.**
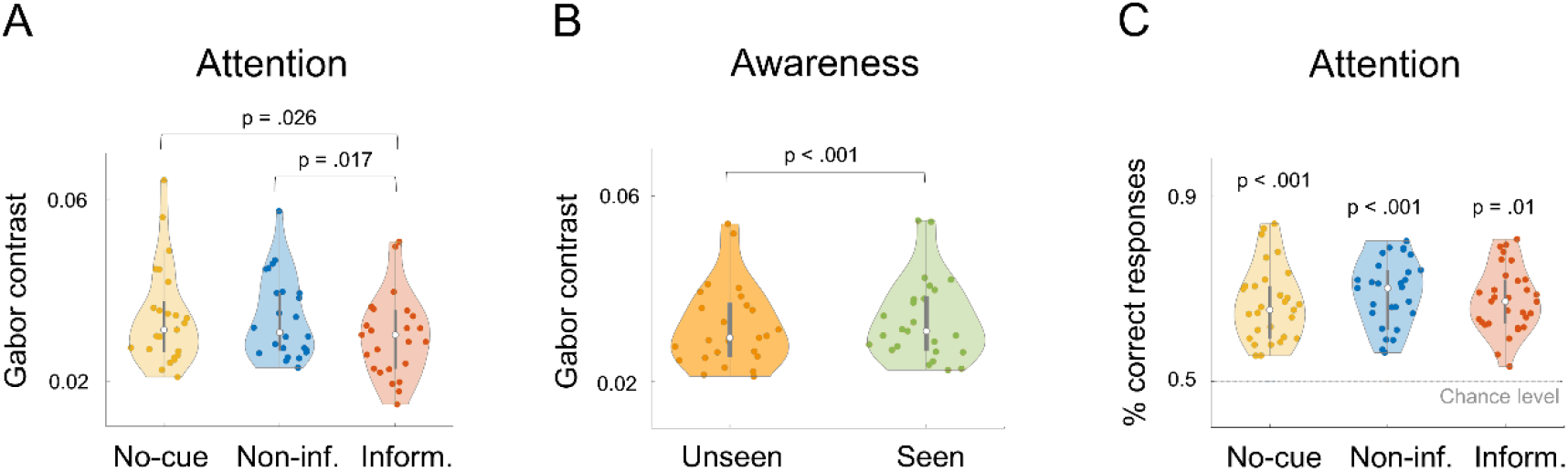
Behavioral results. Violin plots illustrating the results of the behavioral analyses. **A.** Significant main effect of Attention on Gabor contrast. **B.** Significant main effect of Awareness on Gabor contrast. **C.** Proportion of correct responses in the objective question for each task. Participants’ performance was significantly above chance level in all of them. Abbreviations: Non-inf.: non-informative; Inform.: informative.

### Performance in the objective question was above chance level

Performance in the objective forced-choice question was assessed individually for each task by means of t-tests against 0.5. As expected, the proportion of correct responses was above chance level in each of the three tasks: no-cue (t_28_ = 11.24, p < 0.001), non-informative cue (t_27_ = 14.37, p < 0.001), and informative cue (t_30_ = 14.16, p = 0.01) (see Figure 2C).

### Spontaneous posterior alpha amplitude modulates conscious perception

The cluster-based permutation analysis revealed significant differences in pre-stimulus oscillatory amplitude between seen and unseen trials in the no-cue and in the non-informative cue tasks. Specifically, in the no-cue task, we found a negative cluster (p = 0.023) over posterior channels extending from 8 to 12 Hz and from 0.25 s before stimulus onset. In the non-informative cue task, we found a negative cluster (p = 0.017) with similar topography and time-frequency features, although in this case extending from 0.37 s to stimulus onset (Figure 3). In both cases, significant differences resulted from a greater decrease in alpha-band amplitude in seen compared to unseen trials. Critically, we did not find any significant differences in oscillatory amplitude between seen and unseen trials in the informative cue task (p > 0.09).

**Figure 3.**
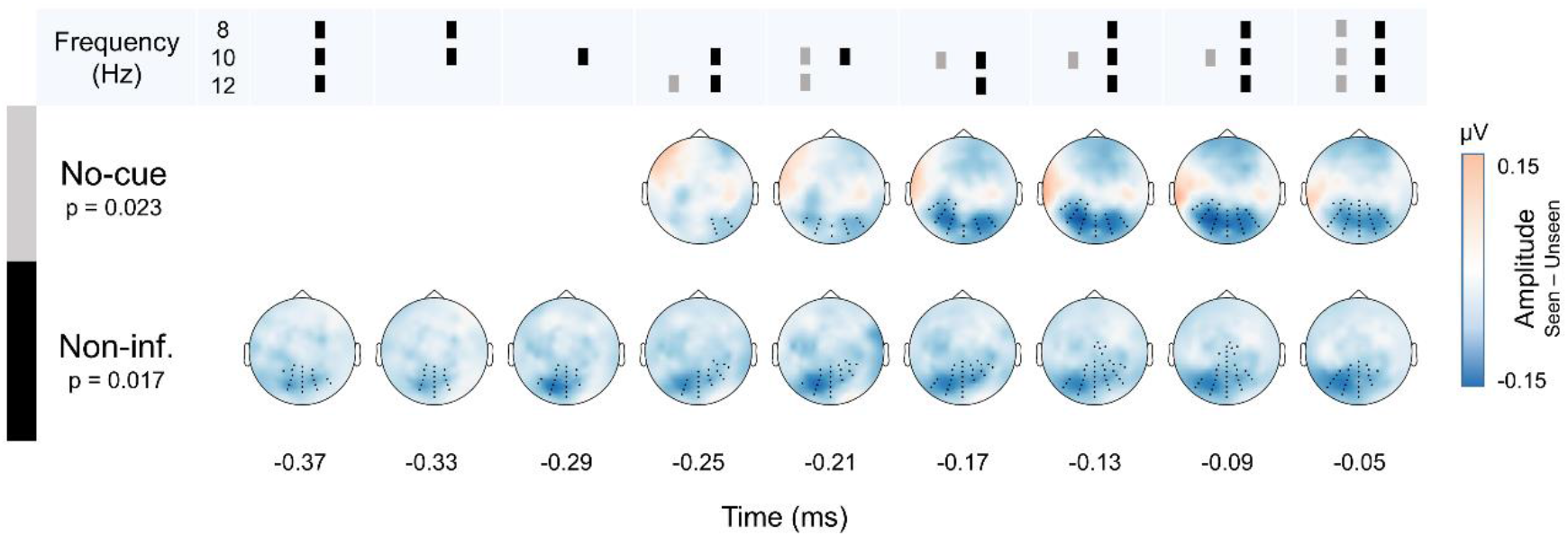
Cluster-based results on pre-stimulus oscillatory amplitude. Grand-average of the amplitude differences between seen and unseen trials. Cluster-based statistics revealed significant differences in the no-cue and the non-informative cue tasks over posterior sites (black dots), extending from 8 to 12 Hz and during 0.25 and 0.37 s previous to stimulus onset, respectively. The upper row indicates the exact frequency bins involved at each time point (grey represents the no-cue task, black represents the non-informative cue task). Abbreviations: Non-inf.: non-informative.

### Absence of phase effects on perceptual outcome

To investigate the role of phase on visual perception, we conducted a cluster-based analysis comparing the empirical POS and the estimated null POS. No significant differences in POS were found in any task. Furthermore, as the pulsed-inhibition theory predicts that pre-stimulus alpha phase would influence perception more when alpha amplitude is high, we repeated the cluster-based permutation tests separately for low and high amplitude data. Results revealed no significant difference in either in the low or high amplitude subset in any of the three tasks.

### Weak and inconsistent effects of pre-stimulus phase in ad-hoc analysis

Additionally, we performed ad-hoc statistical analyses to check whether any weak effects of pre-stimulus phase might appear when relaxing statistical assumptions (e.g., no multiple comparisons correction). However, even in this case, we did not observe any significant effect in trials with low pre-stimulus alpha amplitude (p > 0.12). On the contrary, we did observe significant (uncorrected) phase effects when pre-stimulus alpha amplitude was high. In the no-cue task, we found phase effects between −0.4 and −0.2 s in the beta frequency band (14-18 Hz) (t_28_ = 4.37, puncorr < 0.001). In the non-informative cue task, effects were also observed between −0.4 and −0.2 s, but in a lower frequency range (6-10 Hz) (t_27_ = 2.16, puncorr = 0.039). Finally, in the informative cue task, we detected pre-stimulus phase effects in a later time window between −0.2 and −0.05 s and in the theta band (4-6 Hz) (t_30_ = 2.33, puncorr = 0.020) (Figure 4). The electrodes with the highest POS values varied for the different tasks, with local maxima over central, parietal, and frontal regions.

**Figure 4.**
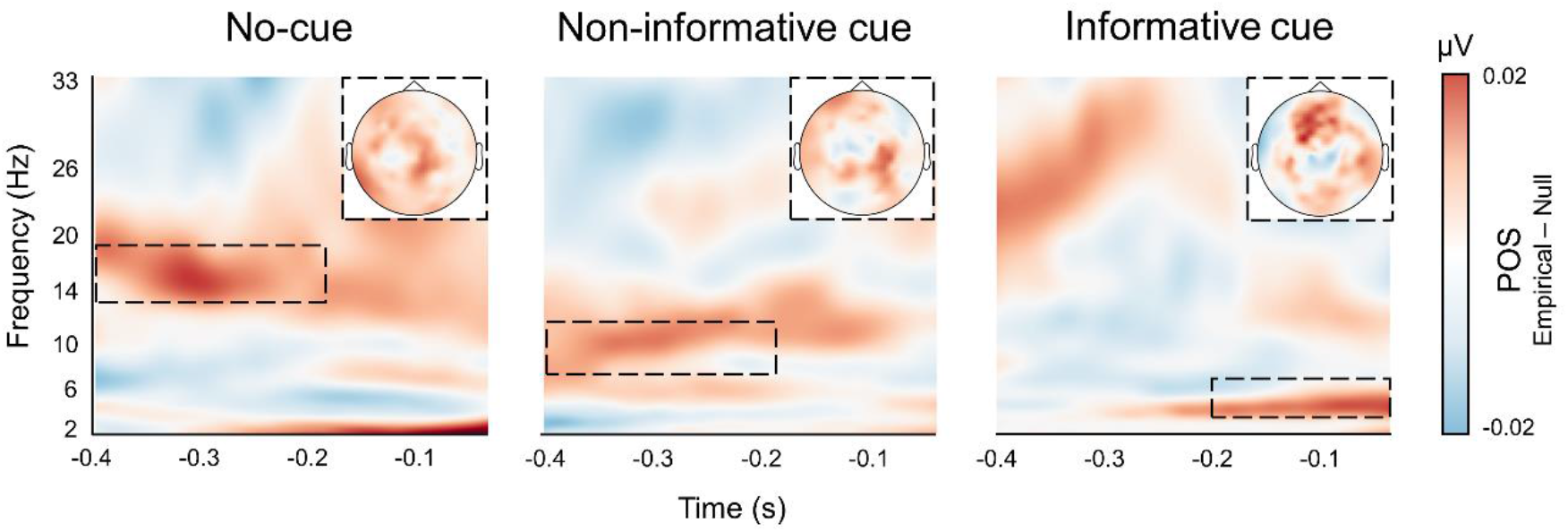
Results of ad-hoc analysis. Ad-hoc pre-stimulus phase effects in trials with high alpha amplitude. Although some uncorrected significant results can be observed, timing, frequencies, and electrodes involved are not consistent across tasks.

### Bayes factor supports amplitude effects and confirms null phase effects

Bayesian analysis was aimed at providing evidence for the null or the alternative hypothesis (H0 and H1, respectively) for both pre-stimulus alpha amplitude and phase. As explained above in the method section, data tested by Bayesian statistics were extracted from a common mask of electrode-time-frequency bins corresponding to the clusters where significant differences in pre-stimulus alpha amplitude appeared in both the no-cue and the non-informative cue tasks (i.e., the only conditions in which significant effects were found). Figure 5A depicts the common channels of the clusters obtained in the two tasks. The size of the dot indicates the number of times each electrode was represented in the cluster, which was used to weight the amplitude/phase values of that electrode. Figure 5B shows the grand-average topography for amplitude (seen vs. unseen) and POS (empirical vs. null) for the three tasks after applying the abovementioned weights to each electrode.

**Figure 5.**
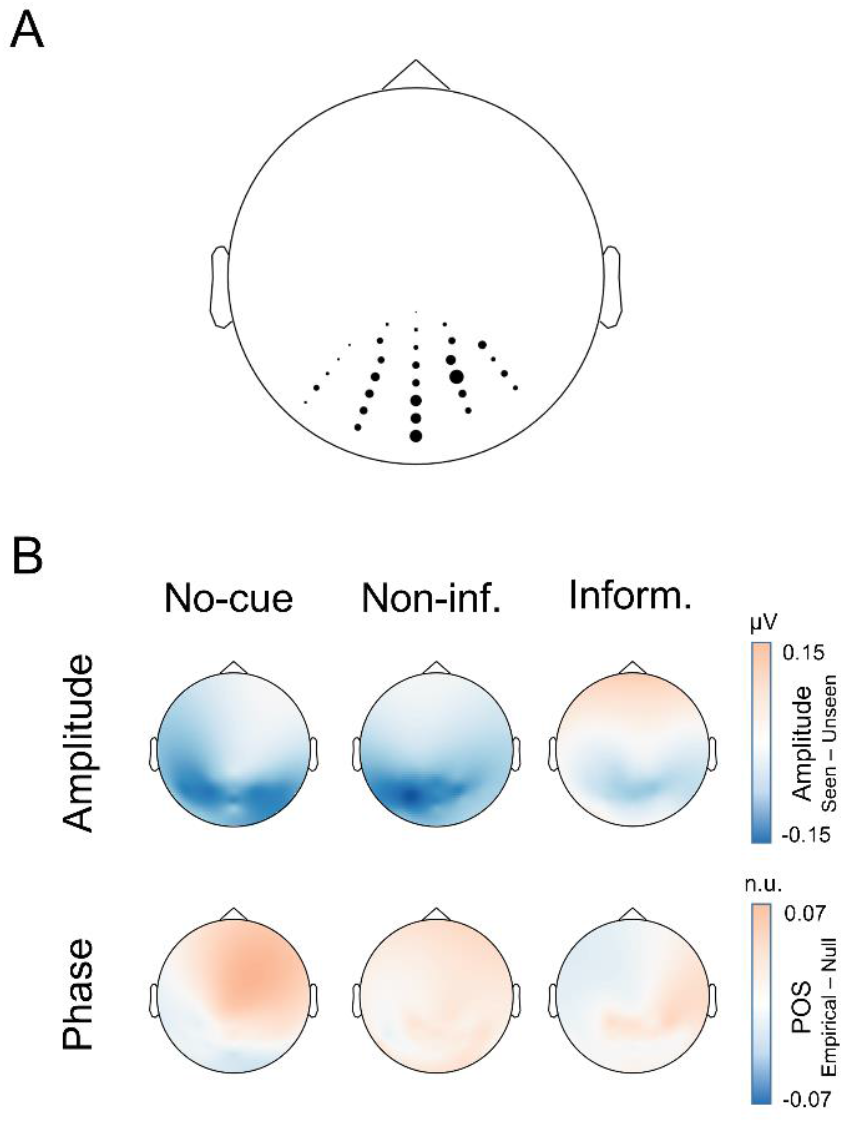
Amplitude and POS values tested by Bayesian analysis. **A.** Common mask derived from the no-cue and the non-informative cue cluster-based amplitude results: 30 weighted posterior electrodes at 10 Hz from −0.25 to −0.05 s before the stimulus onset. **B**. Grand-average amplitude and POS differences for the three tasks after applying the common mask. Abbreviations: Non-inf.: non-informative; Inform.: informative.

The results of Bayesian analysis are summarized in Figure 6A. The analysis of pre-stimulus alpha amplitude revealed strong and moderate evidence in favor of the H1 for the no-cue (BF_10_ = 19.61) and non-informative cue tasks (BF_10_ = 5.24), respectively, whereas the evidence was anecdotal (BF_10_ = 1.10) for the informative cue task. By contrast, alpha phase analysis showed moderate evidence supporting the H0 in both the no-cue (BF_10_ = 0.17) and the informative cue (BF_10_ = 0.19) tasks, while evidence was anecdotal (BF_10_ = 0.62) in the non-informative cue task. Importantly, these results were robust to prior selection, resulting in the same interpretation for a wide range of prior widths (Figure 6B).

**Figure 6.**
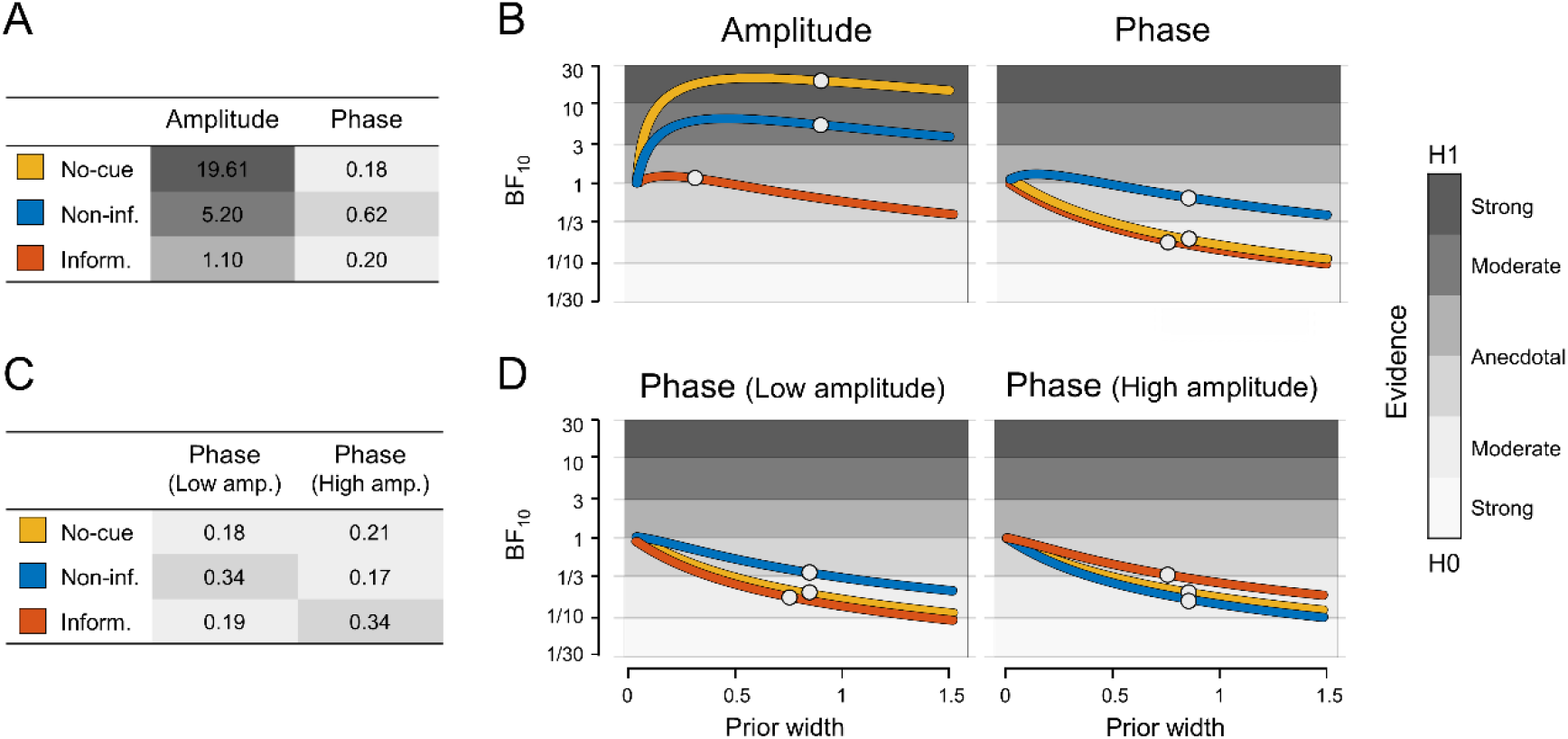
Bayesian results. **A.** Bayes factors for the effect of pre-stimulus alpha amplitude and phase in conscious perception for each task (no-cue, non-informative cue, and informative cue). The strength of the evidence in favor of the H0/H1 is indicated by a gray scale. **B.** Robustness of BF_10_ to different prior widths for pre-stimulus alpha amplitude and phase effects. Priors used are marked by white circles. **C.** BF_10_ for the effect of pre-stimulus alpha phase separately for low and high amplitude trials. **D.** Robustness of BF_10_ to prior specification for the phase effect in low and high amplitude trials. Priors employed are indicated by white circles. Abbreviations: Non-inf.: non-informative; Inform.: informative; Amp.: amplitude.

Finally, we analyzed the evidence in favor of the H0/H1 for alpha phase separately for trials with low and high pre-stimulus amplitude. Taken together, our results show moderate evidence, or a trend towards it, in favor of the H0 for the three tasks and for both low and high amplitude trials (low amplitude trials: no-cue BF_10_ = 0.18, non-informative cue BF_10_ = 0.35, informative cue BF_10_ = 0.19; high amplitude trials: no-cue BF_10_ = 0.21, non-informative cue BF_10_ = 0.17, informative cue BF_10_ = 0.35) (Figure 6C). In all cases the results were stable, supporting the H0 for different prior choices (see Figure 6D). In sum, Bayesian analysis suggests that perceptual outcome is not influenced by the pre-stimulus alpha phase, not even in cases of high alpha amplitude.

## Discussion

This study aimed to shed light onto the role of pre-stimulus alpha-band amplitude and phase in conscious visual perception. Participants were encouraged to detect a near-threshold visual stimulus under different conditions of attentional involvement: no-cue, non-informative cue (50% validity), and informative cue (100% validity). Data analysis was conducted using a data-driven approach, avoiding any a priori selection of parameters. A cluster-based permutation analysis of time-frequency activity revealed a significant decrease of pre-stimulus parieto-occipital alpha amplitude in seen compared to unseen trials, which extended from 8 to 12 Hz and during ~0.3 s prior to stimulus onset. As confirmed by Bayesian statistics, there was moderate to strong evidence for the effect of pre-stimulus alpha amplitude on visual awareness both in the no-cue and the non-informative cue tasks, but not when the cue was 100% informative. On the contrary, we found no evidence of pre-stimulus alpha phase effects for any of the tasks.

### Spontaneous fluctuations in pre-stimulus alpha amplitude influence visual awareness

Our results show a strong effect of pre-stimulus posterior alpha-band amplitude on subsequent perception, but only when alpha was spontaneously fluctuating and not modulated by attention. Thus, in both the no-cue and the non-informative cue task, alpha amplitude was lower during a ~0.3 s pre-stimulus time window when the upcoming peri-threshold stimulus was detected. Our results agree with most previous studies, which consistently find a negative relationship between pre-stimulus alpha power and perceptual awareness (Benwell et al., 2022; Chaumon & Busch, 2014; Ergenoglu et al., 2004; Iemi et al., 2017; Ruzzoli et al., 2019). This effect is commonly attributed to spontaneous fluctuations in cortical excitability (Pfurtscheller, 2001; Romei et al., 2008): when the excitability of visual neurons is higher (and alpha power is lower), forthcoming stimuli are more easily perceived. Combined TMS-EEG studies have added compelling evidence in support of this interpretation by showing that reduced posterior alpha power enhances the likelihood of experiencing TMS-induced illusory phosphenes (Fakche et al., 2022; Romei et al., 2008; Samaha et al., 2017).

Importantly, the deployment of visuospatial attention has also been found to dampen alpha-band power in the visual cortex (Capilla et al., 2014; Sauseng et al., 2005; Thut, 2006; Worden et al., 2000). The question then arises: in the case that alpha oscillations were previously modulated by attention, would they still influence perceptual outcome? Our results show that when pre-stimulus alpha-band oscillations were modulated by a 100% informative cue, alpha amplitude did no longer differentiate between seen and unseen trials. This finding provides insight into the apparent discrepancy of a few earlier studies that failed to find pre-stimulus alpha amplitude effects on perception (Harris et al., 2018; Milton & Pleydell-Pearce, 2016; Wyart & Tallon-Baudry, 2008), since in all these cases predictive cues were employed. We must conclude, then, that only spontaneously fluctuating pre-stimulus alpha-band power has a clear influence on subsequent perception. When alpha oscillations are modulated by attention, perceptual effects disappear, or may at best be weak, as indicated by Bayesian statistics. This might be explained by a ceiling effect. In brief, attention-induced alpha suppression improves overall perception, most likely by raising excitability levels (Capilla et al., 2014; Romei et al., 2008). However, increased excitability might reach a ceiling for both seen and unseen trials and, therefore, pre-stimulus alpha-band amplitude would no longer be decisive for stimulus perception.

The non-informative (50%) cue task was designed to control that the results obtained in the informative (100%) cue task were actually due to the involvement of attention and not to the mere perception of the peripheral cue. Since null pre-stimulus alpha amplitude effects were restricted to the informative cue task, we are confident that they can be attributed to attention. In the same vein, Gabor contrast analysis showed a perceptual advantage only in the informative cue task, indicating that participants effectively deployed anticipatory attention to the cued location. An open question is what would happen with intermediate validity rates between 50 and 100%. It has been nicely demonstrated that as cue validity gradually increases, perceptual performance improves and lateralization of alpha suppression over posterior electrodes becomes more pronounced (Gould et al., 2011). Therefore, one would expect a gradient in the effect of alpha amplitude as a function of validity. However, prior studies using informative cues with validity rates ranging from 60 to 75% have obtained inconsistent results (Capilla et al., 2014; Harris et al., 2018; Milton & Pleydell-Pearce, 2016; Wyart & Tallon-Baudry, 2008), which suggests that once attention is engaged, pre-stimulus alpha power has either no effect or a weak effect on perception.

### Pre-stimulus alpha phase has no influence on visual perception

While the amplitude of spontaneous posterior alpha-band oscillations is consistently found to influence perception, as discussed above, there is more controversy on the role of the alpha phase. According to the rhythmic perception framework (VanRullen, 2016b), there should be an optimal phase for sensory processing within each alpha cycle, while the opposite phase would lead to a poorer perceptual outcome. Thus, pulses of inhibition have been hypothesized to occur every 100 ms, creating windows of optimized processing paced by the phase of alpha oscillations (Klimesch et al., 2007; Mathewson et al., 2011; Mazaheri & Jensen, 2010). Pioneering work (Busch et al., 2009; Mathewson et al., 2009) indeed found that a specific phase of pre-stimulus alpha oscillations facilitated stimulus detection. This effect has been replicated several times (Busch & VanRullen, 2010; Dugue et al., 2011; Fakche et al., 2022; Harris et al., 2018; Samaha et al., 2017; Zazio et al., 2021), although regions (frontal, fronto-central, occipital), time windows (varying within the temporal interval −0.5-0 s) and frequencies involved (4 to 20 Hz) are not consistent. Furthermore, a growing number of studies have reported null effects of alpha phase on visual perception (Benwell et al., 2017, 2022; Chaumon & Busch, 2014; Ruzzoli et al., 2019; Vigué-Guix et al., 2022).

To contribute to this open debate, we extracted pre-stimulus oscillatory phase values per channel-frequency-time point in each of our three tasks, and computed POS values between seen and unseen trials, which should lead to positive values when both conditions show opposite phase clustering (VanRullen, 2016a). Contrary to theoretical predictions, none of the three tasks showed any significant difference in POS values for any frequency band, indicating that there was no pre-stimulus alpha phase associated with better or worse perceptual performance. Bayesian analysis corroborated this finding, by showing moderate evidence for the absence of alpha phase effects in both the no-cue and the informative cue tasks, whereas the evidence for the non-informative cue task was not conclusive. Taken together, our results therefore do not support the view that pre-stimulus alpha phase have an impact on the perception of peri-threshold visual stimuli.

One of the critical factors that may account for the contradictory findings in the literature is the use of different experimental paradigms. In some studies, only one stimulus is presented centrally, while others have used two competing stimuli, which can sometimes be anticipated by a spatial cue. In addition to spatial expectation, the time interval between the cue and the stimulus can be either variable or fixed, which would add a temporal expectation component to the task (see Ruzzoli et al., 2019 for a summary of experimental settings). Thus, anticipatory attention is a key source of divergence between studies. Indeed, a strength of the present study is that the same version of the task, with and without attentional cueing, was applied to the same participants. Although a lack of pre-stimulus alpha phase effect was observed in all tasks, this was slightly more robust when attention was deployed towards the stimulus location.

As previously mentioned, it is a common finding that anticipatory attention induces a suppression of alpha-band power in the cortical regions specialized in processing incoming stimuli (Capilla et al., 2014; Romei et al., 2008). And it is important to note that a pronounced reduction of alpha power may preclude phase effects for two reasons, one methodological and one theoretical. Firstly, low power levels make it technically difficult to obtain a reliable estimate of oscillatory phase (Mathewson et al., 2011; Vigué-Guix et al., 2022). And, secondly, lower alpha power has been hypothesized to provide longer temporal windows for sensory processing (Jensen & Mazaheri, 2010; Mazaheri & Jensen, 2010) which, if alpha suppression is strong enough, could overlap with each other. Hence, attention-induced alpha power reduction might produce a sustained enhancement of cortical excitability, allowing for a consistent level of stimulus processing (Mathewson et al., 2011) and, consequently, an overall perceptual improvement that is not phase-dependent. Critically, in order for alpha suppression to override phase effects, attention should be maintained throughout the entire cue-target interval. For example, the long and variable delay (1-2 s) used by Bush and colleagues (2009) could have resulted in attention-modulated alpha power returning to baseline levels 0.3 s before stimulus onset, which might have facilitated the presence of phase effects thereafter (see Figure 2 in Bush et al., 2009).

### The interaction of pre-stimulus alpha amplitude and phase does not predict perceptual outcome

To examine the potential mediating role of alpha power on the effect of phase, we repeated all phase opposition analyses separately for trials with high and low oscillatory amplitude. However, we still did not find any significant effect. Furthermore, Bayesian analyses revealed moderate evidence for the absence of phase effects in both high and low alpha amplitude trials. Consequently, our results do not provide support for the pulsatile inhibition hypothesis, which predicts that phase opposition between consciously seen and unseen stimuli should become evident during increased levels of alpha power (Klimesch et al., 2007; Mathewson et al., 2011; Mazaheri & Jensen, 2010). It is important to mention that only a few studies have empirically explored the trade-off between alpha amplitude and phase in visual perception, yielding contradictory results. While Mathewson (2009) and Fakche (2022) found evidence supporting theoretical predictions, others have failed to find any interaction between amplitude and phase (Harris et al., 2018; Milton & Pleydell-Pearce, 2016), in line with our findings.

One possibility is that the particular conditions of our experimental design hindered the detection of potential phase effects. We manipulated attention by introducing a 100% valid cue in one of the tasks, whereas in the other two tasks there was no prior information about stimulus location. Although, in the latter case, we assumed no attentional involvement, participants had to perform a relatively demanding task, detecting near-threshold stimuli at either of two peripheral locations without the aid of spatial or temporal cues. Thus, it might be that pre-stimulus alpha oscillations exhibited some degree of modulation by attention in both tasks and, therefore, alpha oscillations did not reach the higher amplitude levels typical of resting. Indeed, previous reports of a positive amplitude-phase interaction have used low attention demanding tasks, such as the detection of a masked stimulus presented centrally after a fixed time interval (Mathewson et al., 2009) or a TMS-induced phosphene perception task (Fakche et al., 2022). This alternative explanation would be in line with the hypothesis that alpha phase plays a particularly important role in the perception of unattended stimuli (see Jensen et al., 2012; although see Harris et al., 2018; Milton & Pleydell-Pearce, 2016).

A second possibility that may explain the lack of a significant alpha amplitude-phase interaction in our study is the lower sensitivity of our statistical analysis to existent but weak effects. We applied cluster-based permutation tests on three-dimensional data (electrode-frequency-time) without making any a priori assumptions about regions or frequency bands of interest, which may have mitigated the multiple comparisons problem. In fact, it is a common practice to focus statistical analysis on either a subset of electrodes (e.g., Busch & VanRullen, 2010; Dugue et al., 2011; Mathewson et al., 2009) or a specific frequency range (e.g., Mathewson et al., 2009; Samaha et al., 2015). To test this possibility, we conducted an additional analysis in which statistical assumptions were strongly relaxed by selecting ad-hoc the time-frequency windows with the highest phase effects. Despite such favorable conditions, we still did not observe any significant effect of pre-stimulus phase in trials with low pre-stimulus alpha amplitude. By contrast, significant (uncorrected) phase effects emerged in high alpha amplitude trials, although they were rather inconsistent, appearing in different time windows and frequency bands (theta, alpha, and beta), and with different topographies in each task. It is important to keep in mind that these results were obtained through a circular method (Kriegeskorte et al., 2009) with the sole purpose of detecting potential weak effects that may not have reached statistical threshold when corrections for multiple comparisons were adequately applied. Therefore, they should not be taken as positive results, but only as possible trends that may help to interpret previous conflicting findings. Indeed, the heterogeneity of our results resembles that observed in the literature, with phase effects reported in different regions, pre-stimulus time windows, and over a wide variety of frequencies ranging from theta to beta. Similar to the conclusions drawn by others (Benwell et al., 2017; Ruzzoli et al., 2019), these results suggest that, in the best-case scenario, pre-stimulus alpha phase has a weak and inconsistent impact on perceptual outcome, compared to the robust effect of pre-stimulus parieto-occipital alpha power.

## Conclusions

Here, we have shown that only the amplitude of spontaneously fluctuating pre-stimulus alpha oscillations have a strong influence on the subsequent perception of near-threshold visual stimuli. This finding reinforces the view that alpha power indexes neural excitability and can therefore be regarded as a filtering mechanism that gates incoming sensory information (Klimesch et al., 2007; Romei et al., 2008). However, the pulsed-inhibition theory rather emphasizes the functional role of the alpha phase, as this can impact perception at much shorter timescales by periodically opening windows for information processing (Jensen & Mazaheri, 2010; Mathewson et al., 2011; Mazaheri & Jensen, 2010). Critically, we did not find any evidence in support for this hypothesis, nor for a potential trade-off between pre-stimulus alpha amplitude and phase. In fact, Bayesian analysis provided moderate evidence in favor of the absence of any phase-related effect. Taken together, the evidence gathered in this study support the view that, unlike pre-stimulus alpha power, the effect of pre-stimulus alpha phase on conscious visual perception is not a robust finding.

Finally, it is worth mentioning that the influence of pre-stimulus alpha amplitude on perceptual outcome is consistently observed over parieto-occipital regions and within the canonical limits of the alpha band (e.g., Chaumon & Busch, 2014; Romei et al., 2008; Ruzzoli et al., 2019). Phase effects are, however, much more heterogeneous. However, the pulsed-inhibition framework implicitly assumes that alpha power and phase effects are two complementary mechanisms arising from the same underlying oscillatory process (Mazaheri & Jensen, 2010). Thus, a convincing demonstration that alpha oscillations control the flow of information processing by modulating both its power and phase should prove that pre-stimulus power and phase effects co-occur in the same region and at the same oscillatory frequency.

## Acknowledgements

This work was supported by FEDER/Ministerio de Ciencia, Innovación y Universidades – Agencia Estatal de Investigación, Spain (grants PGC2018-100682-B-I00 to AC and PC, PID2021-125841NB-I00 to AC, PID2019-111335GA-I00 to CP). The funders had no role in study design, data collection and analysis, decision to publish, or preparation of the manuscript.

## Author Contributions

**María Melcón**: Conceptualization, Data curation, Formal analysis, Investigation, Visualization, Resources, Writing – original draft preparation, Writing – review & editing; **Enrique Stern**: Conceptualization, Data curation, Formal analysis, Investigation, Visualization, Resources, Writing – original draft preparation, Writing – review & editing; **Dominique Kessel:** Formal analysis, Methodology, Writing – review & editing; **Lydia Arana**: Conceptualization, Investigation, Writing – review & editing; **Claudia Poch:** Conceptualization, Funding acquisition, Writing – review & editing; **Pablo Campo**: Conceptualization, Funding acquisition, Project administration, Writing – review & editing; **Almudena Capilla**: Conceptualization, Funding acquisition, Methodology, Project administration, Supervision, Visualization, Writing – original draft preparation, Writing – review & editing.

